# Evaluation of a new probiotic concept for broilers

**DOI:** 10.1101/2020.02.25.964460

**Authors:** S. L. Jørgensen, L. L. Poulsen, M. Bisgaard, H. Christensen

## Abstract

Probiotics were introduced as a spray directly in the hatcher when chickens started to leave the eggs which potentially could reduce the horizontal transmission and colonization with pathogenic bacteria. The single introduction of probiotics could limit the cost compared to multiple introductions with feed and/or water. A mixture of five probiotic strains belonging to *Escherichia coli, Enterococcus faecalis, Lactobacillus agilis* and *Lactobacillus rhamnosus* was tested with two independent flocks of broilers (Ross 308). For each experiment, a comparison was made to an untreated control flock on the same farm. At day 14 of production the probiotic strains were re-isolated from ileum of euthanized chickens. The first week mortality was slightly increased in the probiotic flock (0.42%) compared to the control (0.35%) in experiment 1, however, it was higher in the control flock (1.45%) compared to the probiotic flock (1.12%) in experiment 2. The average weight of chickens that could be slaughtered for consumption was increased by 3.5% in the probiotic flocks compared to the control flocks, resulting in a 1.9% higher total weight of slaughtered chickens in the probiotics treated flocks compared to the control as a mean of the two experiments. The number of condemned animals was within the normal range for the production system and could not directly be related to effects of probiotics. Although one probiotic strain of *E. coli* was isolated from dead animals, the probiotics did not affect the proportion of chickens which died due to *E. coli* during the first week compared to the control.

**Primary audience:** plant managers, veterinarians, nutritionists

## DESCRIPTION OF PROBLEM

Bacterial infections in industrialized chickens can cause high mortality and major economic costs [1, 2]. Bacteria including *Escherichia coli*, enterococci and staphylococci have been reported most frequently to contribute to bacterial disease in poultry, being associated with septicemia, omphalitis, salpingitis and endocarditis [3, 4, 5]. Vertical transmission from the parent hen to the chicks and subsequent horizontal transmission in the hatcher has been considered to be the main route of the infectious agent to spread [5, 6].

The chicken’s first week of live has been found critical and the main bacterial challenge has been *E. coli* [7]. Producers aim to keep first week mortality (**FWM**) below 0.7% [8]. Antibiotics have traditionally been preferred to control FWM. However, due to the development of antibiotic-resistant bacteria, alternative treatment methods have been proposed to reduce bacterial infections.

Newly hatched chicks in industrialized settings can be exposed to the microflora found on the eggshells in the hatchers and bacteria of chicks which may hatch with a vertical transmitted infection [5]. The colonization route of chickens provided the idea to use a new way to deliver probiotics to the chickens. A novel application method (new probiotic concept, **NPC**) has been introduced applying the probiotic bacteria in the hatchers during the hatching proces. The probiotics were sprayed in the hatcher at the time point where 50% of the eggs had hatched. The method relied on the further horizontal transmission of the probiotic strains and colonization of the hatching chicks.

Probiotics have been defined as “a live microorganisms which, when administrated in adequate amounts, confer a health benefit on the host” [9]. Probiotics have been used as prophylactic treatment of industrialized chickens and a wide range of bacterial strains and application methods have been described [10-12]. Several bacterial species have been investigated for a probiotic protential. Most significantly lactobacilli have been shown to have a protective effect against pathogenic bacteria as well as a beneficial effect on the digestive and immune responses in chickens. Additionally, both *Enterococcus faecium* and *Escherichia coli* have been shown to have a similar positive effect in chickens under experimental conditions [13-19].

The application of probiotics in broiler production has been well studied and probiotics in particular have have been demonstrated to have positive effects in the control of pathogenic bacteria, including species of *Campylobacter, Salmonella* as well as *E. coli* [11, 20, 21]. Unfortunately most studies performed and positive results reported have been obtained under experimental conditions and not on farm level.

The objective of this study was to investigate NPC in an industrialized production setting and to test the effect of the NPC on hatchability, FWM, feed conversion ratio (**FCR**), production efficiency factor (**PEF**) and total slaughtered weight.

## MATERIALS AND METHODS

### Approval of the animal experiment

The experiment was approved by The Danish Veterinary and Food Administration, the Ministry of Environment and Food (case no. 2015-29-21-00714).

### Probiotic strains

Strains listed in Table 1 were provided freeze-dried in a powder formulation. Immediately before spraying, the strains were solubilized in deionized water. Whole genome sequencing (**WGS**) was used to characterize the strains genetically and to design strain specific PCR primers for identification [22].

**Table 1.** Probiotic strains and doses of cell forming units (**CFU**) used in the two experiments.

|  |  |  | Dry formulation of probiotics per treatment | Predicted dose | Measured dose in tank when the hatcher was emptied |
| --- | --- | --- | --- | --- | --- |
| Species | Strain | Experiment | g | CFU/ml | CFU/ml |
| *E. coli* | 730-281114-10n | 1 | 20 | $3.2 \times 10^7$ | $5.6 \times 10^6$ |
| *E. coli* | L12 | 2 | 20 | $1.3 \times 10^6$ | ND <sup>1</sup> |
| *E. faecalis* | 336 | 1 | 20 | $3.7 \times 10^5$ | $7.6 \times 10^7$ |
| *L. agilis* | 268a | 1 | 20 | $2.9 \times 10^8$ | $2.2 \times 10^7$ |
| | | 2 | 20 | $2.9 \times 10^8$ | $2.2 \times 10^7$ |
| *L. rhamnosus* | SACCO SP1 | 1 | 10 | $2.1 \times 10^8$ | ND |
| | | 2 | 10 | $2.1 \times 10^8$ | ND |
| ND <sup>1</sup>, not detected |  |  |  |  |  |

### Distribution of probiotics directly in the hatcher

The hatchers were of model Petersime (Zulte, Belgium) [25]. The spray devise to deliver the probiotic spray was designed as an independent unit that was placed in the hatcher at the same time that it was loaded with fertile eggs [26]. A dose of probiotics was approximately 5×10^8^ cell forming units (**CFU**) for each chick and was delivered in 1.0 ml deionized water. After spraying, a sample of the probiotic solution was collected for counting of CFU (Table 1).

### Farm

The NPC was tested in a conventional Danish broiler production farm. Experiments 1 and 2 were carried out on the same farm in two similar buildings with respect to size, feed, watering system, ventilation and temperature controls. In the first experiment 15,400 Ross 308 chicks were inserted in both the probiotic treated and the control group. In the second experiment 15,000 chicks were included each group (Table 2). In experiment 1, the probiotic treated chicks were inserted in house 1, and the control chicks in house 2. In the second experiment, chicks treated with probiotics were inserted in house 2 and house 1 left for the control flock. The farm performed a thinning after 34 days where 5,000 and 4,500 broilers were removed for slaughter from each house, respectively from the two experiments followed by a final delivery to slaughter at day 38. The effect of the probiotics was measured by FCR [27], FWM and PEF [27], as well as total and average weight of birds at slaughter.

**Table 2.**
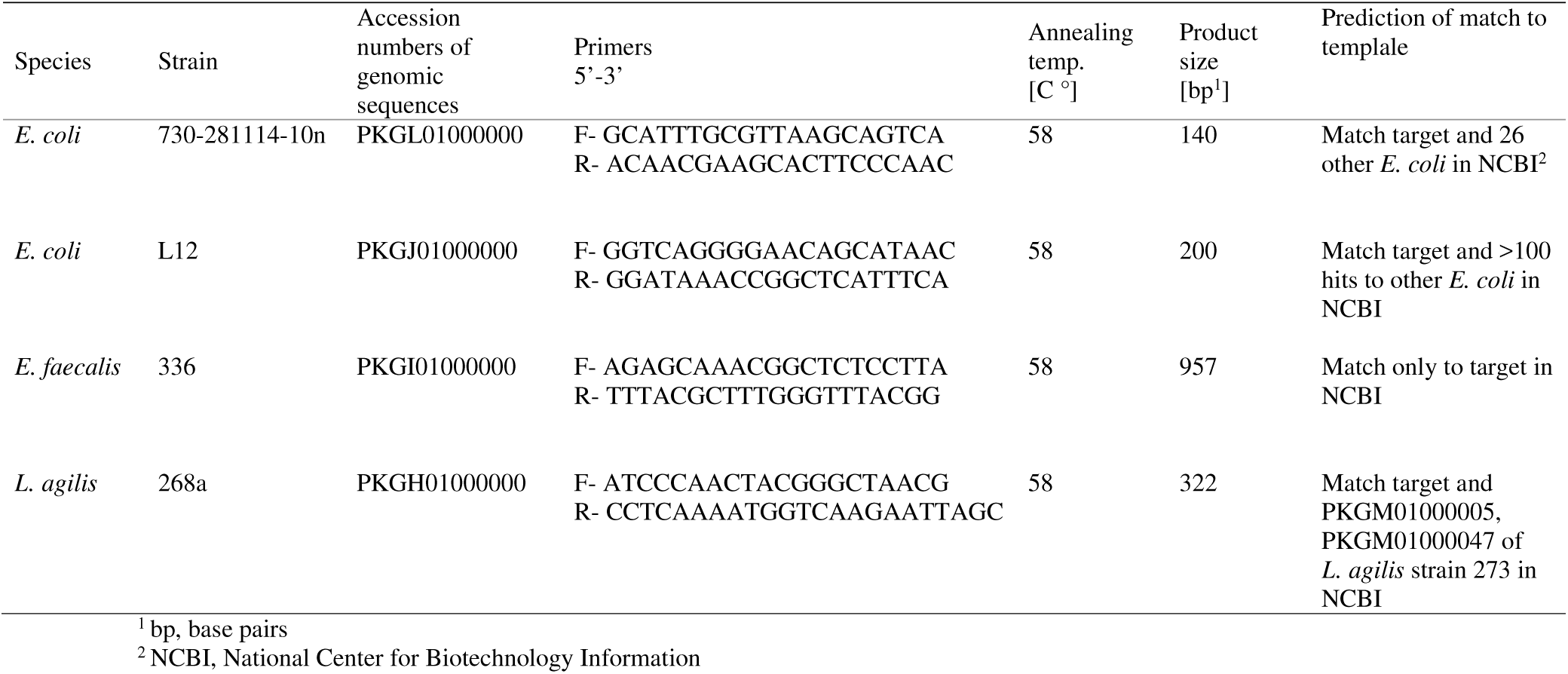
Genomic accession numbers of strains, PCR primers and PCR conditions used to detect probiotic strains.

At day 14, 25 birds from the treatment and control flocks were euthanized by cervical dislocation and contents of ileum (n=25) and cecum (n=25) were collected and tested for the presense of the probiotic strains by culturing and PCR. Dead chickens were collected throughout the production period and immediately stored at – 20 °C.

### Probiotic strain recovery and identification

The content of the ileum from the 25 euthanized birds (approximately 1 g wet weight per bird) were frozen in glycerol (15% vol/vol final conc.) and stored at - 80 °C until further analysis. Samples were plated on Man de Rogosa, Sharpe broth [29] for recovery of lactobacilli and blood agar plates (**BA**) (5% calf blood in blood agar base [30] for recovery of *Enterococcus faecalis* and *E. coli* and incubated overnight at 37 °C. From all BA and MRS agar plates, five presumptive *E. coli, E. faecalis* and *L. agilis* colonies were isolated per plate and pooled before suspension in in 100 μL sterile filtered ultraclean water [31] for preparation of DNA templates. The samples were boiled for 10 min and centrifuged at 14,000 ×g for 5 minutes. The supernatant was used as DNA template for PCR. PCR identification of the probiotic samples was made, using strain-specific primers (Table 2). To confirm the presence of the probiotic *E. faecalis* and *L. agilis* strain, boiling lysates were prepared in 85 μL sterile filtered ultraclean water with 5 μl lysozyme (10 μg/ml) and 10 μl lysis buffer (200 mM Tris HCl, 500 mM KCl, pH 8.4) at 37 °C for 20 min. Hereafter, 4 μl protein kinase K was added and digested for 60 min at 56 °C. The samples were centrifuged at 15,000 ×g for 5 min at 5 °C, the supernatant was transferred to a new tube and used for DNA PCR template. For samples found positive by PCR, single colonies from the primary plates were confirmed positive by the strain-specific PCR tests. Five single colonies were used to prepare DNA template from each sample.

### Post mortem *examination of dead chickens*

The birds collected underwent a full *post mortem* examination including bacteriological sampling upon indication when vascular disturbances, exudations or enlargement of the liver was present. The yolk sac and/or liver were sampled with a sterile cotton swab after sterilising the tissue with a hot burning iron

### *Bacteriology of* E. coli *and* E. faecalis *isolates collected during* post mortem *examination*

Bacteriological samples collected during *post mortem* examinations were plated on BA and grown aerobically overnight at 37 °C. From plates showing abundant and pure growth of presumptive *E. coli* colonies (circular with entire margins, low convex, medium size, non-transparent with a light greyish colour on BA) or *E. faecalis* colonies (smooth, circular colonies with entire margins, low covex and small size with a white to greyish colour), one single colony per sample was sub-cultured in Brain Heart Infusion Broth (**BHI**) [32] and stored in 15% glycerol at −80°C. All isolates were tested with the strain-specific PCRs to verify whether the isolates were the *E. coli* and *E. faecalis* used as probiotics. In addition, all isolates which tested positive in the strain-specific PCR were typed by Pulsed-Field-Gel-Electrophoresis (PFGE) for comparison to the probiotic *E. coli* and *E. faecalis* (Table 1). The PFGE protocol for *E. coli* was used in a modified version [33]. The restriction enzyme *Xbal* was used [34]. The PFGE protocol for *E. faecalis* was described by Jørgensen et al. [35].

### Statistical methods

Fischer’s exact test was used for the analysis of data from GraphPad Prism5 [36] with a level of significance of 95%.

## RESULTS AND DISCUSSION

### Delivery of spray in the hatcher

The spray device and delivery of the spray was initially tested without eggs in a hatcher with paper in the trays. Deionized water with Aviblue [37] as trace dye was used. Over 50% of the trays received the dye after a full spray event. Aerosols will also be taken up by the chicks and a higher proportion than 50% of chickens will probably receive the probiotics with a full hatcher.

The hatchability was not affected by the NPC treatment. The humidity increased from 29.3 to 30.8% during the 7 s activation of the spray device, however, the spray device did not affect the automatic humidity control in the hatcher.

### Survival of probiotic bacteria in the hatcher

During the 24 h that the probiotic cultures were left in the hatcher at 37 °C, the CFU decreased with approximately one order of magnitude (Table 1).

### *FWM and* post mortem *examiniation*

In experiment 1, the FWM was 0.42 and 0.35% in the treated and the control house, respectively. With FWM below 0.7%, the two flocks can be regarded as normal performing in regard of FWM [8]. In the second experiment FWM was 1.45 and 1.12% in the control and treated house, respectively. For this experiment the FWM may be regarded as higher than expected according to the broiler management manual [8]. However, the broiler breeders of the flock in experiment 2 were 54 weeks compared to 37 weeks in the first experiment which could explain the higher mortality. A total of 233 chickens dead on their own were collected for *post mortem* and bacteriological analysis. The most prominent lesions for chicken that died during the first week were yolk sac infections, sepsis, peritonitis, pericarditis and perihepatitis. For the first experiment, only 26 and 16% of the chickens collected as dead on their own during the first week from the probiotics and control flock, respectively allowed the isolation of *E. coli* in pure growth which could indicate an effect of the probiotic treatment, however, for the second experiment the similar figures were 73 and 95% for the probiotics and control flock, respectively [38] indicating that the probiotics did not affect the proportion of chickens which died due to *E. coli* during the first week.

The strains isolated during *post mortem* analysis were investigated with the strain specific PCR methods developed for the probiotic strains and were furthermore investigated by PFGE. In Experiment 1, none of the *E. coli* isolated from diseased chickens of the control group were identified as the probiotic *E. coli* strain by the strain-specific PCR. However, four of the *E. coli* isolated from the probiotic treated flock were positive for the strain-specific PCR of strain 730-281114-10n and showed identical PFGE banding patterns to the strain delivered in the inoculum. Three isolates were obtained from yolk sac infections and the fourth from sepsis. Furthermore, the two *E. faecalis* strains isolated tested positive in the strain-specific PCR for probiotic *E. faecalis* strain 336 and showed identical banding patterns by PFGE analysis compared to the probiotic strain. The two isolates were obtained from yolk sac infection and sepsis, respectively. The probiotic *E. coli* strain 730-281114-10n and *E. faecalis* strain 336 were excluded from experiment 2 based on the negative results.

### Re-isolation of probiotic bacteria from 14 day old chicks

In experiment 1, the probiotic *E. faecalis* strain 336 and *E. coli* strain 730-281114-10n were identified by PCR in 4 and 8% of collected ileum samples, respectively. The probiotic *L. agilis* strain 268a was identified by PCR in 20 and 60% of the collected cecum and ileum samples, respectively.

In experiment 2, the probiotic *E. coli* strain L12 was identified by PCR in 8% of the collected samples, while *L. agilis* strain 268a was identified by PCR in 10 and 35% of the collected cecum and ileum samples, respectively. Two isolates of *L. agilis* from ileum were subsequently analysed by WGS and SNP analysis to confirm the identity to strain 268a of *L. agilis*. The two sequenceses only differed from the 268a genome by a single SNP and the isolates were concluded to be identical to the original strain.

In previous studies performed in experimental animal facilities it was demonstrated that the probiotic bacteria used in these two field experiments could be re-isolated from the majority of the treated chickens (data not shown). The intestinal flora will probably not change dramatically with time and the probiotic strains may be transient bacteria colonizing the chickens for the first weeks. In the current study the probiotic strain added with NPC were present during the first period of the production period where the chickens show the highest susceptibility to infections by pathogenic bacteria. The commercial strain SP1 was not tested for survival in the current study but has, previously been shown to survive for 30 days at 4°C in emmer beverage [39].

### Total weight and condemnation at slaughter

The total mortality included dead animals in production, animals that died during transport and condemned animals. The total mortality was significantly higher in the probiotic flocks compared to the control flocks in both experiments (Table 3). The higher mortality in the probiotic treated flocks was balanced out by an increased in mean weight in the probiotic flocks. The mean weight of chickens accepted for consumption was 3.3 and 3.7% higher in the probiotic flocks compared to the control in experiment 1 and 2, respectively. The final slaughtered weigth accepted for consumption was higher in the probiotic groups compared to the control flocks in both experiments by 1.7 and 2.1% for experiment 1 and 2, respectively (Table 3).

**Table 3.**
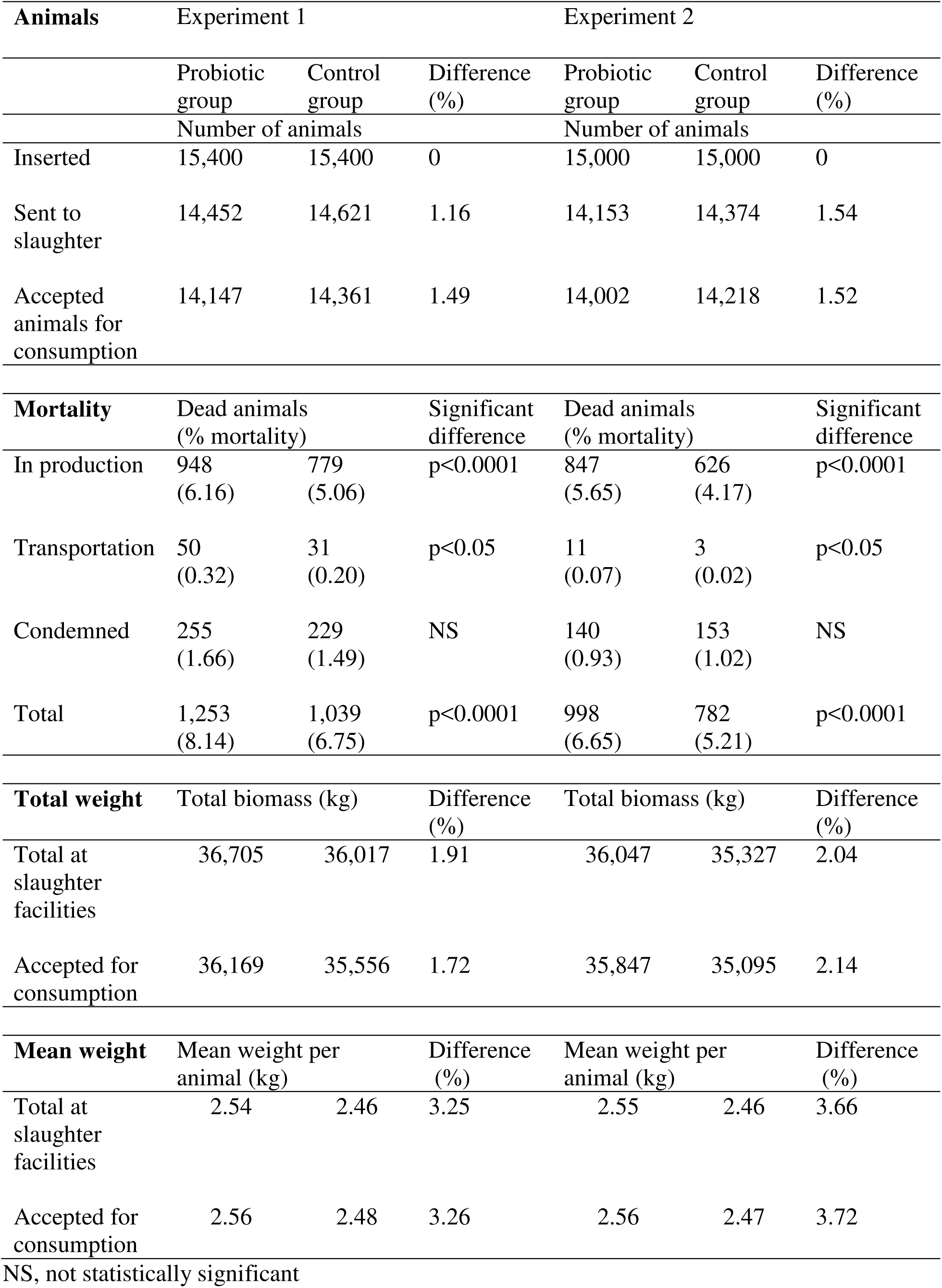
Production parameters for experiment 1 and 2.

| **Animals** | Experiment 1 |  |  | Experiment 2 |  |  |
| --- | --- | --- | --- | --- | --- | --- |
|  | Probiotic group | Control group | Difference (%) | Probiotic group | Control group | Difference (%) |
|  | Number of animals |  |  | Number of animals |  |  |
| Inserted | 15,400 | 15,400 | 0 | 15,000 | 15,000 | 0 |
| Sent to slaughter | 14,452 | 14,621 | 1.16 | 14,153 | 14,374 | 1.54 |
| Accepted animals for consumption | 14,147 | 14,361 | 1.49 | 14,002 | 14,218 | 1.52 |
| **Mortality** | Dead animals (% mortality) |  | Significant difference | Dead animals (% mortality) |  | Significant difference |
| In production | 948<br>(6.16) | 779<br>(5.06) | p<0.0001 | 847<br>(5.65) | 626<br>(4.17) | p<0.0001 |
| Transportation | 50<br>(0.32) | 31<br>(0.20) | p<0.05 | 11<br>(0.07) | 3<br>(0.02) | p<0.05 |
| Condemned | 255<br>(1.66) | 229<br>(1.49) | NS | 140<br>(0.93) | 153<br>(1.02) | NS |
| Total | 1,253<br>(8.14) | 1,039<br>(6.75) | p<0.0001 | 998<br>(6.65) | 782<br>(5.21) | p<0.0001 |
| **Total weight** | Total biomass (kg) |  | Difference (%) | Total biomass (kg) |  | Difference (%) |
| Total at slaughter facilities | 36,705 | 36,017 | 1.91 | 36,047 | 35,327 | 2.04 |
| Accepted for consumption | 36,169 | 35,556 | 1.72 | 35,847 | 35,095 | 2.14 |
| **Mean weight** | Mean weight per animal (kg) |  | Difference (%) | Mean weight per animal (kg) |  | Difference (%) |
| Total at slaughter facilities | 2.54 | 2.46 | 3.25 | 2.55 | 2.46 | 3.66 |
| Accepted for consumption | 2.56 | 2.48 | 3.26 | 2.56 | 2.47 | 3.72 |

2 NS, not statistically significant

The number of condemned animals was higher in the probiotic treated flock (1.66%) compared to the control flock (1.49%) in the first experiment, while the number of condemned animals was higher in the control flock (1.02%) compared to the probiotic flock (0.93%) in the second experiment, neither were significant (Table 4).

**Table 4.** Percentage of animals in the different condemnation categories.

| Experiment | House | Treatment | Skin discoloration | Irregular color/odor/consistency | Ascites | Stunted growth | Entrails remaining | Lack of bleeding off |
| --- | --- | --- | --- | --- | --- | --- | --- | --- |
| 1 | 1 | Probiotics | 14.5 | 54.9 | 14.5 | 4.71 | 5.88 | 5.49 |
| 1 | 2 | Control | 19.7 | 53.7 | 7.86 | 7.42 | 2.18 | 6.99 |
| 2 | 2 | Probiotics | 20.7 | 29.3 | 21.4 | 6.43 | 17.9 | 4.29 |
| 2 | 1 | Control | 14.4 | 35.9 | 15.7 | 1.31 | 26.1 | 6.54 |

The number of condemned animals was significantly higher in the first experiment compared to the second in both the probiotic houses and the control houses (p<0.001). The proportion of condemned animals in the two houses were comparable to the three additional houses of the farm (data not shown) and to the normal in Danish production [40]. The higher condemnation in the first experiment compared to the second was mainly due to a significantly higher number chickens condemned due to irregular color/smell/consistency (p<0.001) (Table 4). This type of condemnation could not be directly related to infectious disease and was therefore not considered to have any association with the NPC treatment. Condemnations related to ascites were higher for the probiotics treatments compared to controls for both experiments, however, this condition predominaly has been associated with non-infectious causes. The number of animals condemned due to skin discoloration was not consistently related to the treatment with probiotics.

#### FCR and PEF

The PEF [28] index includes body weight, mortality and FCR [28] and can be used to evaluate the efficacy of the broiler productionon where a high value indicates an effective production. At the day of slaughter in the first experiment, the PEFs were 351 and 365 for the control flock and probiotic treated flock, respectively (Table 5). In the second experiment the PEFs were 379 and 375 for the control and probiotic treated flocks, respectively. However, PEF values include weight, mortality and FCR of the animals sent to the slaughter facilities. When calculating the adjusted FCR and adjusted PEF for the animals accepted for consumption, the FCR per accepted animal was poorer and the adjusted PEF was overall lower for all groups compared to the control. However, the adjusted PEF was higher in the probiotic flocks in both experiments (Table 5).

**Table 5.** Production efficiency for experiments 1 and 2.

|  | **Experiment 1** |  | **Experiment 2** |  |
| --- | --- | --- | --- | --- |
|  | **Probiotic** | **Control** | **Probiotic** | **Control** |
| Feed Conversion Ratio<br>(**FCR**) | 1.61 | 1.64 | 1.56 | 1.58 |
| Production Efficiency Factor<br>(**PEF**) | 365 | 351 | 375 | 379 |
| Adjusted FCR for animals<br>accepted for consumption | 1.70 | 1.73 | 1.66 | 1.64 |
| Adjusted PEF for animals<br>accepted for consumption | 355 | 343 | 372 | 367 |

## CONCLUSIONS AND APPLICATIONS

1. The test of the new probiotic concept under production conditions showed that the probiotic bacteria sprayed in the hatcher during the hatching process did not harm the chicks. However, in the first experiment, one probiotic strain of *E. coli* and one of *E. faecalis* was isolated from dead chicken in production and these strains were not included in the second experiment.
2. A higher total mortality was observed in the probiotic treatments compared to the control and the reasons need further examination. However, FWM including *E. coli* related infections or condemnations were not higher in the probiotics treated flocks compared to the control.
3. The mean weight of chickens and the total slaughtered weight accepted for consumption was higher in the probiotic flocks compared to the control in both experiments.
4. The higher weight of animals in the probiotic treatment compared to the control was reflected in a higher PEF in the probiotic treated flocks compared to the control flocks but only when animals available for consumption were considered in the calculations.
5. Further investigations with the new probiotic concept are suggested to be done to test the persistence of the probiotics in the flocks and to test the concept under different management conditions.

## Acknowledgements

The NPC project was financed by GUDP (Grøn udviklings og demonstrationsprogram), administrated by NaturErhvervsstyrelsen. The project was a collaboration between Danhatch A/S, Sacco and University of Copenhagen. We thank Katrine Aagaard for excellent technical assistance.

